# DNA structure change induced by guanosine radicals – a theoretical and spectroscopic study

**DOI:** 10.1101/312413

**Authors:** J. Kobierski, E. Lipiec

## Abstract

Proton radiation therapy is one of the newest and most promising methods used in modern oncology. Nonetheless, the dissemination of that method may result challenging. This is partially due to the fact that the mechanism of DNA damage induced by protons, which is one of the ways proton radiation interacts with tissues, has not been fully understood yet. It is well known that ionizing radiation especially ions such as protons may directly damage DNA but it also causes the formation of radicals, which may lead to even more serious damage of the DNA sugar-backbone than direct interaction with charged ion. In this article we focus on the influence of guanine radicals on the DNA structure, namely the conformation and stability of the DNA strand. We present the theoretical results of the optimization calculations of DNA structures with guanine radical-adenine pairs as well as calculated Raman spectra. By combining theoretical calculations with the experimental spectra we were able to explain molecular modifications of the DNA sugar-backbone affected by guanine radicals formed upon radiation exposure, which lead to spectral changes between spectra of control and irradiated DNA. Thus we established a pathway of the formation of DNA damage caused by protons.

## Introduction

Proton therapy has been developing for over 60 years now. As far back as 1946 Robert Wilson proposed the use of accelerator–produced beams of protons to treat deep–seated tumors in the human body (1). Nine years later the first patient was treated with proton beams in the Lawrence Berkeley Laboratory (2). First ocular melanoma was exposed to protons in the mid–1970s (3). According to statistics of Particle Therapy Co–Operative Group, up to the present proton therapy has been used to treat 131240 patients in 46 centers worldwide (4). However, in comparison to other radiation therapies an adoption of proton therapy is fairly slow. One of the main reasons is the lack of knowledge about proton interaction with biomolecules.

DNA is the most sensitive biomolecule for radiation exposure (5, 6). A damaged lipid or protein molecule can be easily replaced, whereas damaged DNA must be repaired. Ionizing radiation can cause various types of DNA damage such as DNA strand cross–links, DNA–protein cross–links, Single Strand Breaks (SSBs), base damage, sugar damage and the most dangerous form Double Strand Break (DSBs). In eukaryotes DSBs are critical lesions that can lead to cell death (7).

DNA damage was investigated by plethora experimental and theoretical approaches. Comet assay, which is practically an electrophoresis of single cells, is commonly used in order to detect DNA strand breaks (8). Mass spectrometry and liquid chromatography are efficient tools used to investigate the chemical structure of damaged DNA (9, 10). All these experimental methods can affect DNA samples, leading to changes in their structure due to required DNA degradation or via application of chemical substances and complex preparation procedures (11). Raman spectroscopy, in turn, is a label free and non–invasive analytical technique, which can explore chemical structure of DNA damaged by radiation. However, Raman spectrum of such a complex molecule as DNA contains a lot of overlapping spectral features related to DNA functional groups. Therefore, an interpretation of spectral changes related with exposure to radiation can be problematic. In order to assign Raman bands and find the background of spectral changes related with irradiation we have proposed a combination of experimental and theoretical approach.

In the experimental Raman spectra an appearance of strong bands related to stretching of C=O and NH_2_ banding mode from cytosine and guanine were observed after proton exposure. Therefore, we focused on guanine-cytosine pair and molecular changes, that could be a consequence of interaction with the guanine radical – the most frequently formed radical in the DNA structure. We optimized structures with radicals (not their products) in order investigate direct influence of guanine radicals on the local DNA structure and predict following molecular damage.

Many molecular processes can be simulated by the quantum mechanical calculations. Until now theoretical calculations, in particular Density Functional Theory (DFT), have been successfully used to present models explaining mechanism of DNA damage, including radiation induced DNA damage. Recently these calculations have been used to elucidate DNA strand breaks after irradiation, where ionization of the sugar phosphate backbone by water radicals •OH or direct damage by ionizing particles were considered as the most likely mechanism of DNA lesions (12–14).

Part of research focused on the processes occurring within or around the nitrogen base pairs. Shimuzu et al. used DFT calculations in order to investigate dehydrogenation reaction between GC and AT base pairs and •OH radical, which can be produced upon irradiation of DNA (15). Zhang and Eriksson focused, in turn, on intramolecular radical cross–link reactions in deoxyribonucleosides induced by •OH (16).

Guanine, among all the DNA bases, has the lowest ionization potential, which means that it is the main oxidation center in DNA (17, 18). For this reason, irradiation of DNA leads to the formation of a large number of oxidized guanine (G^·+^). The oxidation of guanine itself and the formation of guanine neutral radicals (•G) in the GC pair have been thoroughly investigated. Studies include breaking hydrogen bonds in the GC pair (19, 20), the formation of cross–linked structures between DNA base pairs (21), or the formation of neighboring bases structures in single strained DNA (22). However, due to limited computing performance, each time only isolated base systems were taken into consideration. Up to the present no results on effects of the interaction disturbance between the nitrogen bases on the DNA backbone or the global double strained DNA structure have been reported.

It is known from previous research that oxidized guanine loses protons either from N1 or N2 site, forming neutral •G_N1_ and •G_N2_ radicals, respectively (23). In this work we present calculated Raman spectra for DNA with the normally bonded GC pair, the DNA with •G_N1_C and •G_N2_C pairs, and the DNA with the pair of cytosine and guanine with hydrogens detached from both N1 and N2 site •G_N1_,_N2_C (Figure 1) along with experimental Raman spectra of undamaged DNA and DNA irradiated by the dose of 4000 protons. By combining the experimental results with theoretical calculations we were able to establish the types of lesions produced by both proton radiation in DNA. Calculations were performed using Density Functional Theory (DFT). Apply of DFT allowed to obtain high precision for the GC pair. The rest of the system was treated classically using the ONIOM hybrid model. The ONIOM method was previously used for Raman spectra calculations of radiation-induced damaged DNA. Simulated spectra were in good agreement with the experimental data (24). Additionally, this model was successfully applied to study DNA interactions with metal complexes (25–27), chloridazon herbicide (28), and proflavine (29).

**Figure 1.**
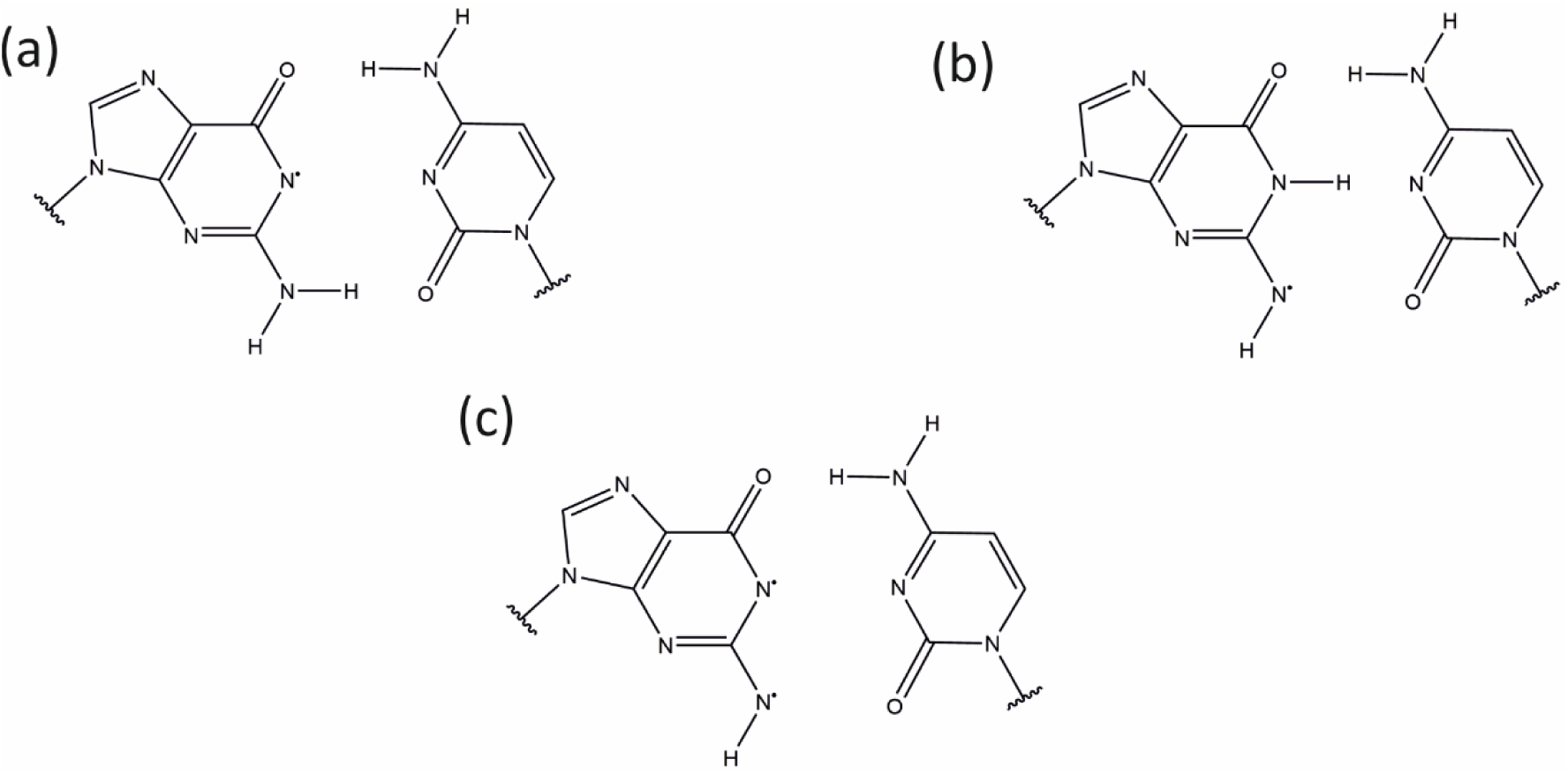
Chemical structural formula of •G_N1_C pair (a), •G_N2_C pair (b), and •G_N1_,_N2_C pair (c)

## Materials and methods

### Experimental procedures

#### DNA irradiation

DNA plasmid pUC–18 (SIGMA Aldrich) was irradiated with 2 MeV protons. A droplet of 2 μl of DNA (concentration 0.5 μg/μl) dissolved in buffer (0.5 mg/ml in 10 mM Tris–HCl with 1 mM EDTA, pH 8.0) was deposited as a thin film on Mylar Foil (Goodfellow Cambridge Limited, Huntington, UK). The Cracow microbeam from the Van de Graaff accelerator was used as the proton source. Samples were irradiated in raster mode with the beam step comparable to the beam diameter ~20μm. The beam current was set to 4000 protons per sec. Two different protons numbers per raster step were applied: 400 and 4000 protons. After irradiation the samples were transferred onto aluminum coated (80 nm) Petri dishes (35 nm diameter) and measured using the Raman microscope.

#### Raman spectroscopy and data processing

Experimental Raman spectra were recorded using a WiTec Confocal Raman Microscope α-300 R equipped with Nd-Yag 532 nm (green) laser with a CCD camera cooled to −60°C. 50x Zeiss objective was applied. Data were collected in the spectral range of 3700 cm^−1^ – 200 cm^−1^. The acquisition time was 5–20 second per spectrum, depending on signal to noise ratio. The spectral resolution was 2 cm^−1^. Raman spectra were processed using Opus 6.5 software, which included smoothing (number of smoothing points: 13) and baseline correction (rubber-band correction, number of baseline points: 16, number of iterations 2–4). In these experiments spectra were vector normalized in the spectral range of 1800 cm^−1^ – 250 cm^−1^ because of the variation in sample thickness.

### Calculation procedures

The DNA oligomer with the AGAGCTCT sequence in B conformation was chosen as a starting structure for optimization. Initial coordinates of atoms were taken from Nucleic Acid Database [http://ndbserver.rutgers.edu]. Geometry optimization and Raman activity calculations for undamaged and radiation–induce damaged (with neutral guanine radical in guanine–cytosine pair) DNA structures were carried out using GAUSSIAN 09 software package. Frequency calculations were performed based on previously optimized structures. All calculations were carried out using ONIOM model, with Becke–Half–and–Half–LYP Density Functional Theory method (BHandHLYP) (30) using 6–311++G(d,p) basis functions for one guanine-cytosine pair with adjacent deoxyribose molecules and a phosphate groups, and Universal Force Field model for additional 7 base pairs. BHandHLYP, a hybrid exchange–correlation functional, was reported as the best performer for the hydrogen–bonded base pairs systems (21). The system was optimized using default integration grid, default integral cutoffs and EDIIS+CDIIS convergence algorithm without damping.

Density Functional Theory methods are known to overestimate vibrational frequencies, which is caused by anharmonicity effects in the theoretical treatment. To avoid this factor and to eliminate known systematic errors resulting from fragmentary incorporation of electron correlation and the use of finite basis sets frequencies obtained from vibrational analysis were scaled by 0.967 factor.

Then, the theoretical Raman intensities were calculated according to the formula (31):

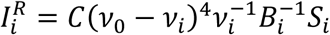

where *C* is a constant given in arbitrary units, *S*_*i*_ and *ν*_*i*_ are the Raman scattering activity and the frequency of the normal mode *Q*_*i*_, respectively. *v_0_* is the frequency of the laser excitation line. In this work this frequency was taken 18797 cm^−1^, what corresponds to 532 nm, a wavelength emitted by Nd:YAG laser used in experiments. *B*_*i*_ is a temperature factor which accounts for the intensity contribution of excited vibrational states and is represented by the Boltzmann distribution:

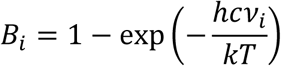

where *h*, *k*, *c* are Planck constant, Boltzmann constant, and speed of light, respectively. *T* stands for temperature, which was assumed 273.15 K.

## Results and Discussion

### DNA

Figure 2 shows a comparison of an experimental spectrum collected from control DNA and theoretical spectrum, calculated for molecular system presented in Figure 3. Signal from DNA bases in both spectra is dominated by ν(C=O) and δ(NH_2_) of cytosine and guanine, base stacking vibration at ~1668 cm^−1^ (24, 32–34), in plane ring vibration of cytosine and guanine as well as δ(NH_2_) in spectral range from 1570 cm^−1^ to 1630 cm^−1^ and ν(C=N) of cytosine together with deformational motion C_2_H_2_ functional group (34, 35). Signal from the DNA backbone involves asymmetric and symmetric stretching of phosphate at 1220 cm^−1^ – 1250 cm^−1^, and at 1040 cm^−1^ – 1110 cm^−1^ respectively and symmetric stretching vibration of O-P-O in the spectral range from 840 cm^−1^ to 770 cm^−1^ (24, 34, 36, 37).

**Figure 2.**
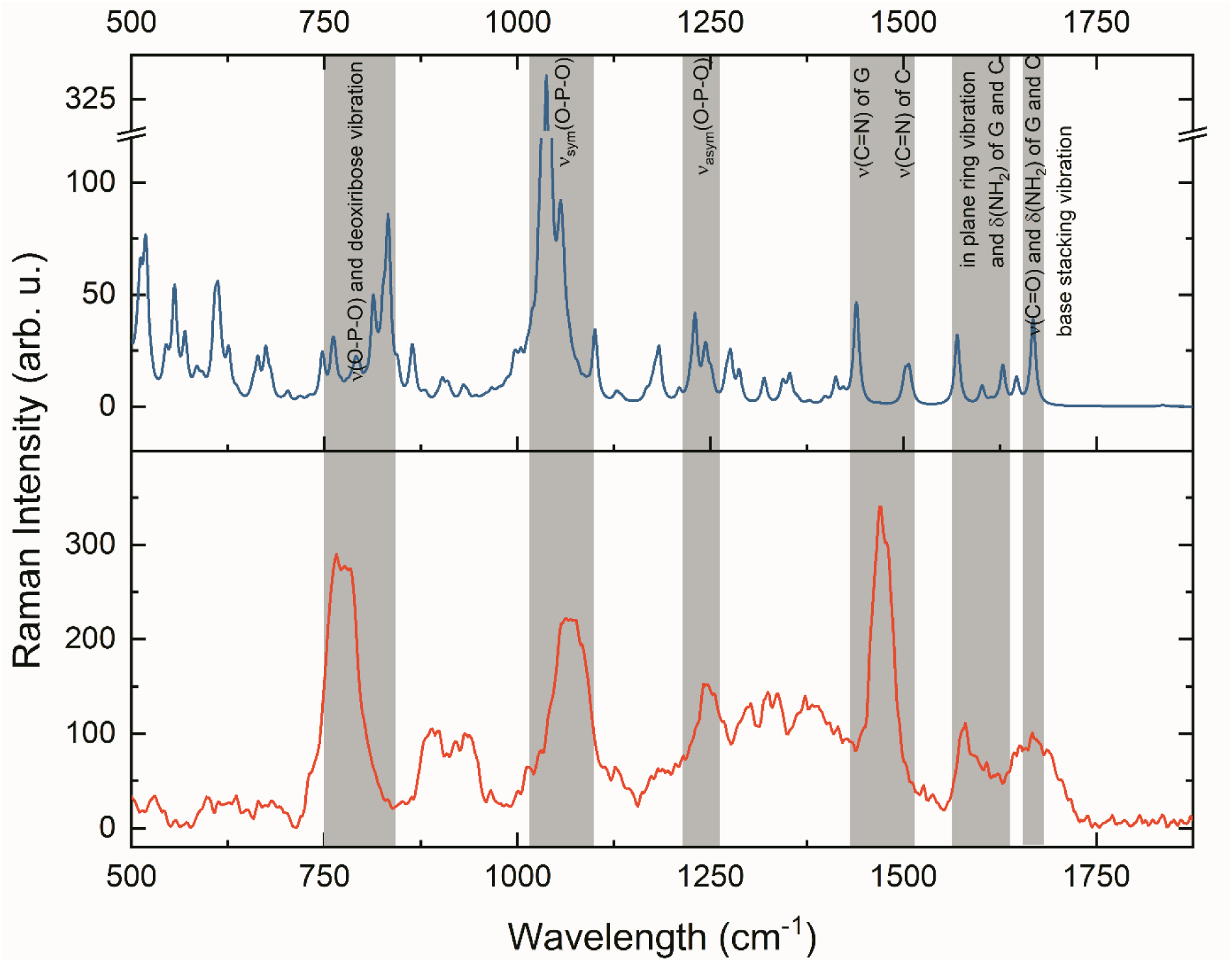
Experimental Raman spectrum along (red) with calculated Raman spectrum of undamaged DNA (blue)

**Figure 3.**
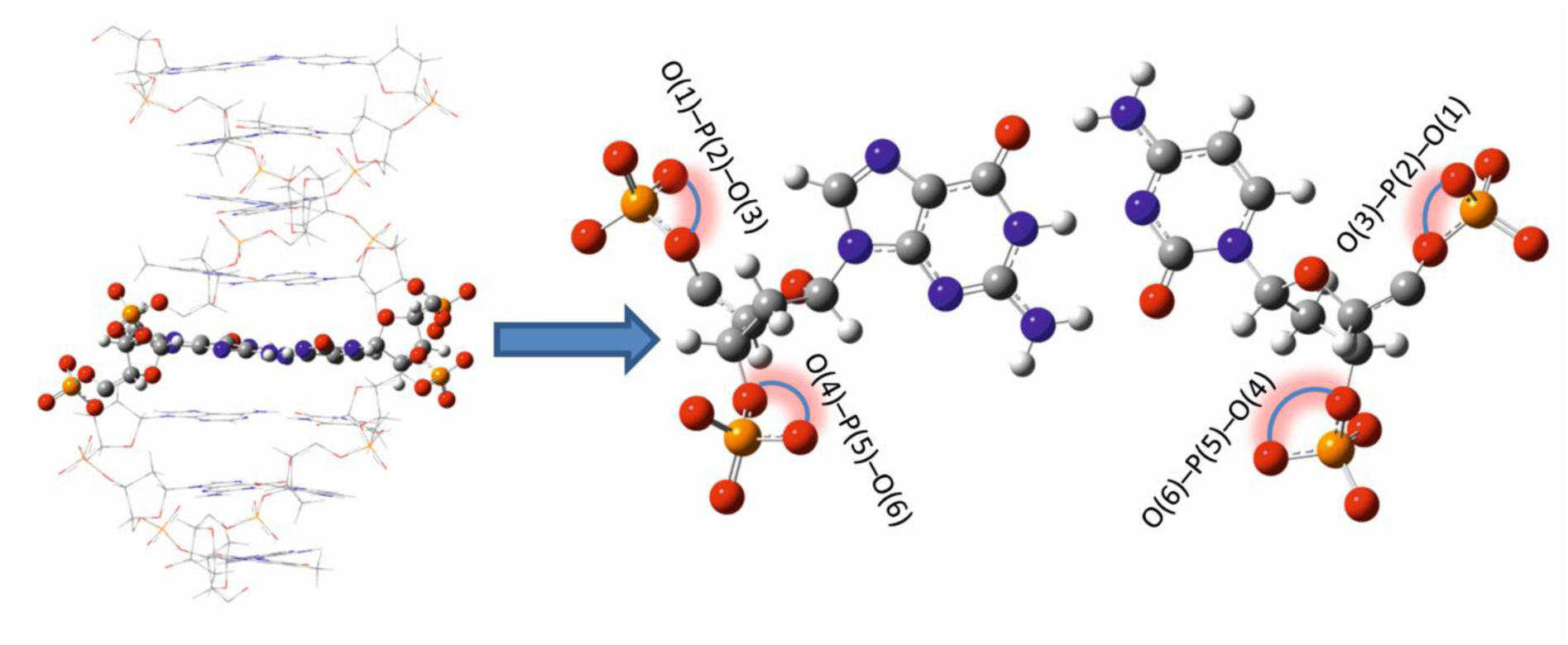
Schematic model of the theoretical molecular system of DNA (left) in which one GC pair (right) was treated quantum mechanically.

Spectra of damaged DNA (irradiated with the dose of 400 and 4000 protons) are presented in Figure 4. Spectra collected from irradiated DNA indicate an appearance of strong bands related to C=O stretching and NH_2_ bending from cytosine and guanine. These bands were not observed after exposure to photons, as described in our previous work (24). Therefore we have assumed that these changes in Raman spectra are related either with the direct radiation damage caused by proton or are possibly related to interaction with guanine radicals, which are the most frequently formed radicals in the DNA structure. Hence, we have tested the hypothesis, that guanine radicals may locally destabilize DNA structure and cause permanent damage, that could be detected in Raman spectroscopy. The lifetime of guanine radicals is in the range of ms. However, they can permanently modify the molecular structure of the DNA backbone (38). We have optimized three DNA structures with various guanine radicals: •G_N1_C, •G_N2_C, •G_N1_,_N2_C and we deeply analyzed their influence on the local molecular structure of the DNA double strand. We generated Raman spectra and compared them with the experimental data to check whether we are able to detect these local molecular changes by Raman spectroscopy.

**Figure 4.**
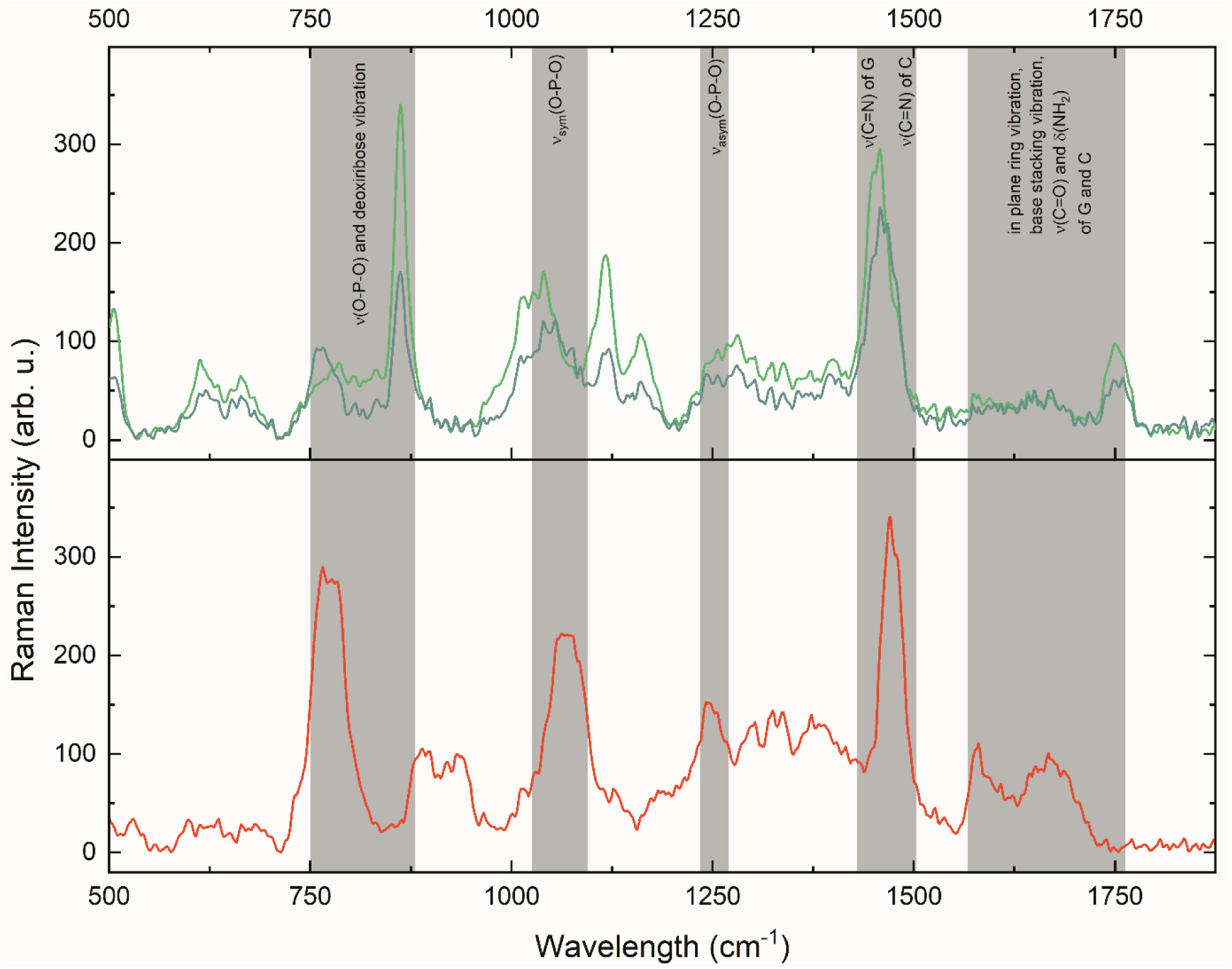
The comparison of experimental spectra of undamaged DNA (red) and DNA irradiated with the dose of 400 protons (cyan) 4000 protons (lime green)

The experimental spectra for 400 and 4000 protons do not differ much from each other as presented in Fig. 4, therefore in the further analysis we focus only on the spectrum for a larger dose, namely 4000 protons. Detailed band wavenumbers and their assignment are reported in Table 1. Experimental data show that bands interpreted as a double bond stretching vibration ν(C=O), scissoring vibration δ(NH_2_) and stretching vibration ν(C=N) of bases cytosine and guanine at 1667 cm^−1^ and 1580 cm^−1^ are shifted after irradiation into 1751 cm^−1^. Band interpreted as symmetric stretching ν_sym_(O–P–O) at 1069 cm^−1^ is splitted after irradiation, whereas stretching vibration band ν(O–P–O) observed at 772 cm^−1^ in spectrum of control DNA is shifted into higher wavelength of 862 cm^−1^ in spectra of irradiated DNA.

**Table 1.** Detailed Band assignments for experimental and calculated Raman spectra od undamaged and irradiated DNA(24, 32-37)

| DNA control | Simulated DNA | Irradiated DNA | DNA with $\bullet\text{G}_{\text{N1}}\text{C}$ | DNA with $\bullet\text{G}_{\text{N2}}\text{C}$ | DNA with $\bullet\text{G}_{\text{N2,N2}}\text{C}$ | Assignment |
| --- | --- | --- | --- | --- | --- | --- |
| 1667 | 1668 | 1751 | 1782 | 1788 | 1782 | $\nu(\text{C}=\text{O})$, $\delta(\text{NH}_2)$ of C, base stacking vibration |
| | | | 1720 | 1760 | 1756 | $\nu(\text{C}=\text{O})$ of G |
| | | | | — | — | $\delta(\text{NH}_2)$ of G |
| 1570-1630 | 1627 | | 1680 | 1708 | 1720 | $\delta(\text{NH}_2)$, in plane ring vibration of C |
| | 1570 | | 1636 | — | — | $\delta(\text{NH}_2)$ of G |
|  |  |  |  | 1602 | 1712 | in plane ring vibration of G |
| 1430-1580 | 1506 | 1460 | — | — | 1574 | $\nu(\text{C}=\text{N})$ of C, def. $\text{C}_2\text{H}_2$ |
| | 1439 | | — | — | 1538 | $\nu(\text{C}=\text{N})$ of G |
| 1220-1250 | 1230 | 1250 | — | — | — | $\nu_{\text{asym}}(\text{O}-\text{P}-\text{O})$ |
| 1040-1110 | 1037 | 1117 | — | 1070 | 1121 | $\nu_{\text{sym}}(\text{O}-\text{P}-\text{O})$ |
|  |  | 1040 |  |  |  |  |
| 772 | 833 | 862 | 857 | 853 | — | $\nu(\text{O}-\text{P}-\text{O})$ and deoxyribose vibration |

All spectral changes of O-P-O bands are associated with conformational changes resulting in shortening or increasing the distance between base pairs. Conformation change require a modification of O-P-O bond length and O-P-O bond angle (Figure 3), which can be directly observed in Raman spectrum as shifts of O–P–O stretching vibrations related bands (36, 37). Therefore, here we are presenting O-P-O bond lengths and O-P-O bond angles obtained for each optimized DNA structure in order to monitor local conformational change induced by DNA damage. The averaged O–P–O bond angle in optimized structure of control DNA was equal to 108.85° ± 0.16°, whereas the averaged P–O bond length was 1.5826 ± 0.0047 Å (Table 2 and Figure 5). Detailed bond length and angles are presented in Table 3. This distance is consistent with the results of crystallographic experiments by Langridge et al. which was 1.60 Å (39).

**Figure 5.**
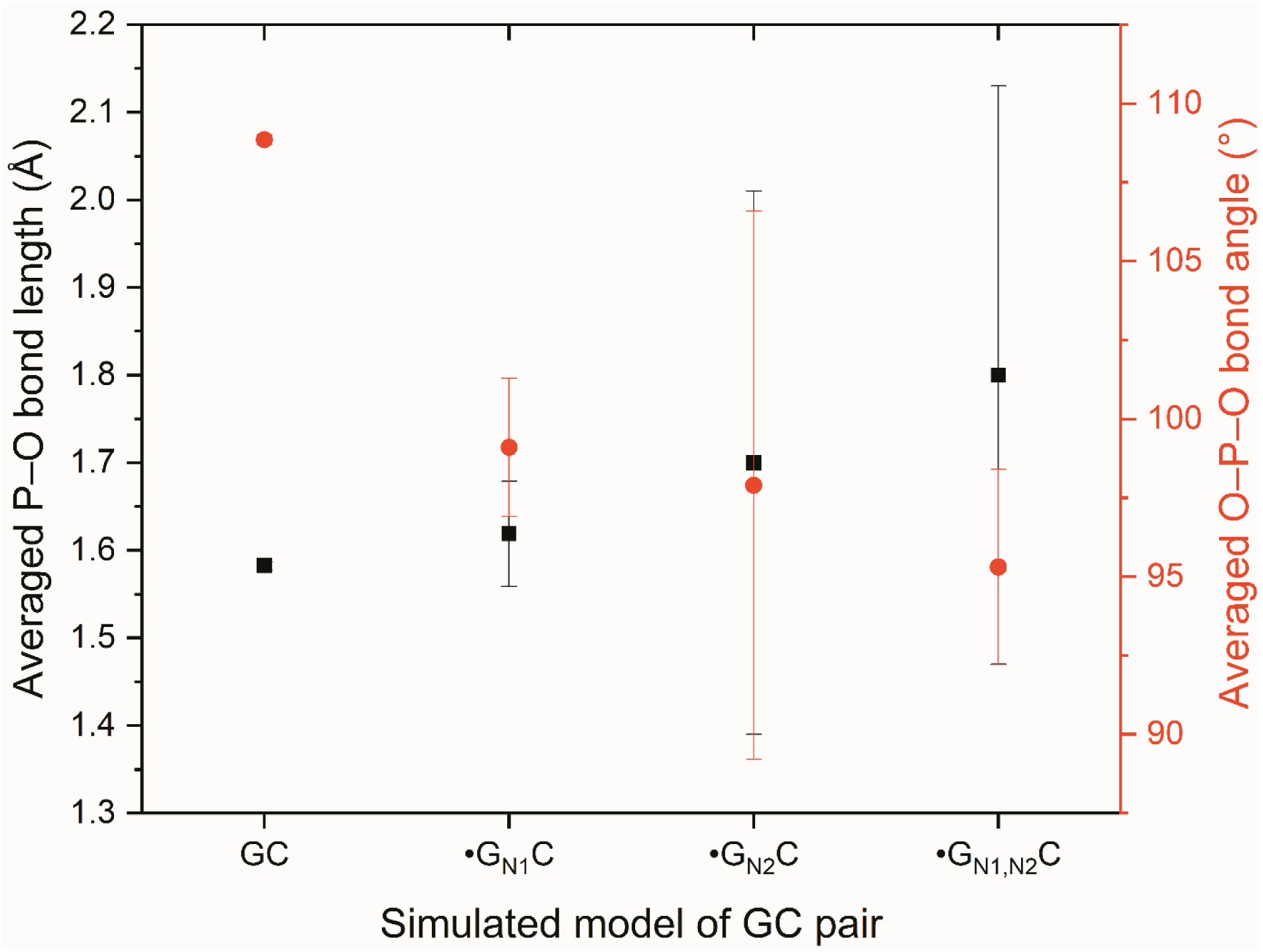
Averaged P-O bond lengths (black squares) and O-P-O bond angles (red circles) obtained in calculations for unchanged DNA and DNA with neutral guanine radicals in GC pair

**Table 2.** Averaged O-P-O bond angles and P-O bond lengths obtained in calculations for unchanged DNA and DNA with neutral guanine radicals in GC pair

| | DNA control | $\bullet\text{G}_{\text{N1}}\text{C}$ | $\bullet\text{G}_{\text{N2}}\text{C}$ | $\bullet\text{G}_{\text{N2,N2}}\text{C}$ |
| --- | --- | --- | --- | --- |
| O–P–O bond angles | $108.85^\circ \pm 0.16^\circ$ | $99.1^\circ \pm 2.2^\circ$ | $97.9^\circ \pm 8.7^\circ$ | $95.3^\circ \pm 3.1^\circ$ |
| P–O bond lengths | $1.5826 \pm 0.0047$ | $1.619 \pm 0.060$ | $1.70 \pm 0.31$ | $1.80 \pm 0.33$ |

**Table 3.** P-O bond lengths and O-P-O bond angles obtained in calculations for unchanged DNA and DNA with neutral guanine radicals in GC pair. For atoms labels explanation see Figure 3, 7, 9, and 11

|  |  | O(1)–P(2)<br>distance (Å) | P(2)–O(3)<br>distance (Å) | O(1)–P(2)–<br>O(3) angle | O(4)–P(5)<br>distance (Å) | P(5)–O(6)<br>distance (Å) | O(4)–P(5)–<br>O(6) angle |
| --- | --- | --- | --- | --- | --- | --- | --- |
| DNA<br>control | G site | 1.59 | 1.58 | 109.09° | 1.59 | 1.58 | 108.66° |
|  | C site | 1.58 | 1.59 | 108.81° | 1.58 | 1.59 | 108.85° |
| •G<sub>N1</sub>C | G site | 1.57 | 1.64 | 98.47° | 1.58 | 1.57 | 102.64° |
|  | C site | 1.70 | 1.55 | 98.36° | 1.62 | 1.72 | 96.81° |
| •G<sub>N2</sub>C | G site | 1.57 | 1.58 | 99.53° | 1.58 | 1.57 | 103.09° |
|  | C site | 1.58 | 2.53 | 83.41° | 1.58 | 1.59 | 105.74° |
| •G<sub>N2,N2</sub><br>C | G site | 1.58 | 2.08 | 92.34° | 2.57 | 1.57 | 92.50° |
|  | C site | 1.68 | 1.55 | 99.87° | 1.63 | 1.72 | 96.31° |

### •G_N1_C

Figure 6 shows experimental spectra of irradiated DNA and calculated spectrum of DNA with •G_N1_ radical in GC pair: •G_N1_C (see Figure 7 for a schematic model). It can be seen, that modes which correspond to stretching vibration ν(C=O) and scissoring vibration δ(NH_2_) are significantly separated and shifted into 1782 cm^−1^ and 1720 cm^−1^ of cytosine and 1680 cm^−1^ and 1636 cm^−1^ of guanine. The band ν_asym_(O–P–O) and ν_sym_(O–P–O) are not significant in this calculation. Experimental peak at 862 cm^−1^ corresponds to calculated mode at 857 cm^−1^ of deoxyribose vibration and stretching vibration band ν(O–P–O). The P–O bond length in this structure is 1.619 ± 0.060 Å, while the angle of O–P–O band is smaller than in unchanged DNA and it equals 99.1°± 2.2°. In this optimized structure we observed the shift of the bases with respect to each other and formation of “unnatural” base pair with only two hydrogen bonds, like in Reynisson work (20).

**Figure 6.**
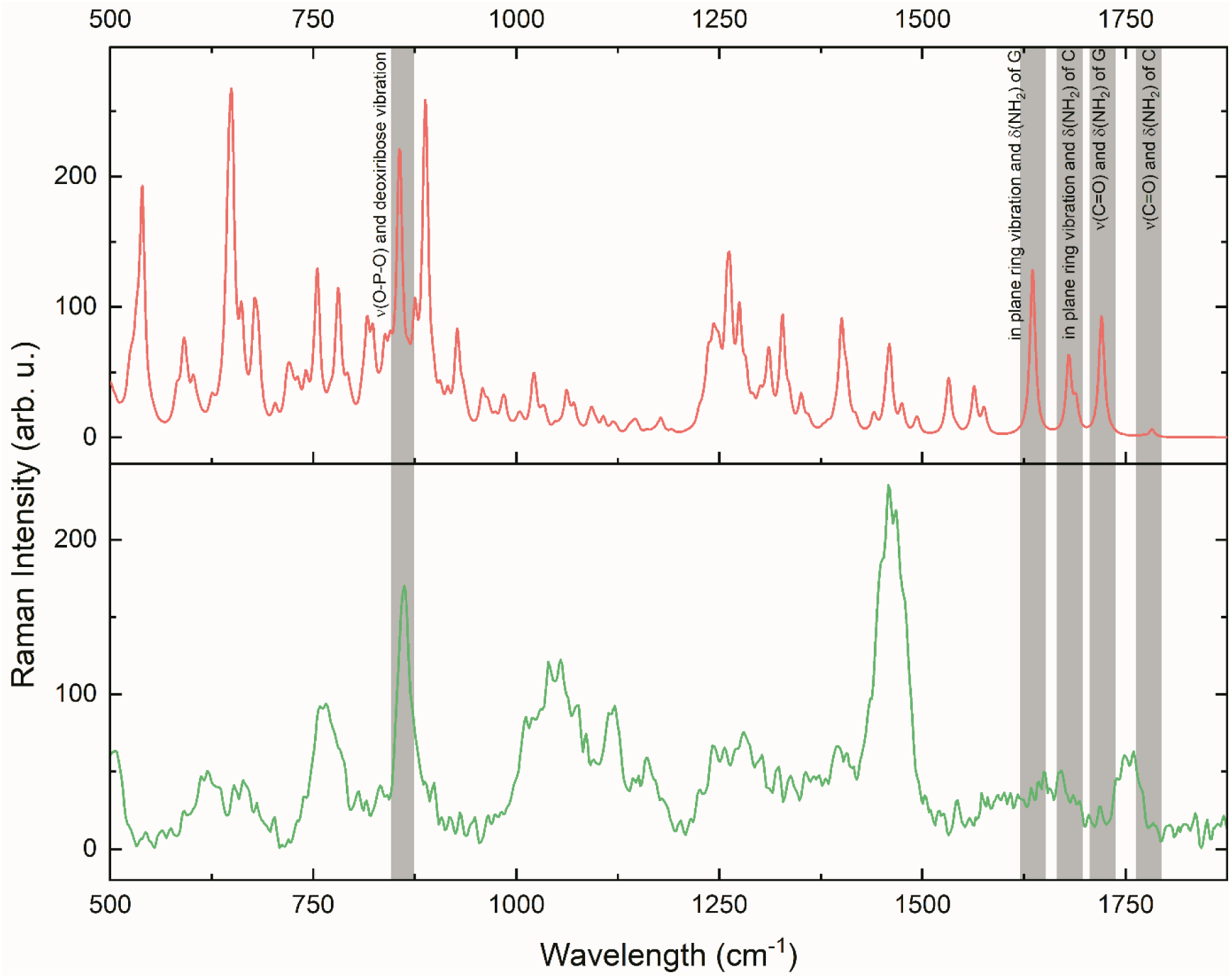
The comparison of spectra of DNA irradiated with the dose of 4000 protons (lime green) with calculated Raman spectrum of damaged DNA with •G_N1_C pair (light red)

**Figure 7.**
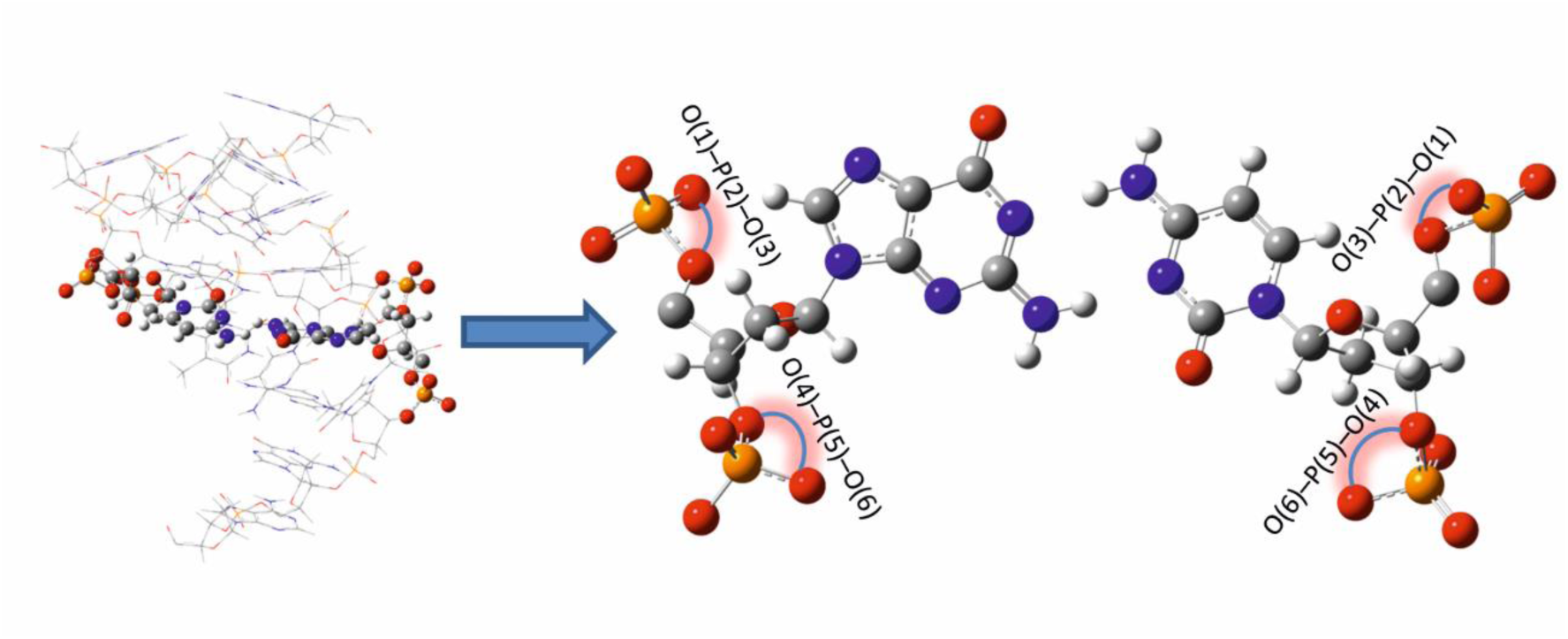
Schematic model of the theoretical molecular system of DNA (left) with •G_N1_C pair (right) treated quantum mechanically

### •G_N2_C

Spectrum calculated for DNA with •G_N2_C pair compared with experimental spectra of irradiated DNA are presented in Figure 8. A schematic model of the optimized structure is shown in Figure 9. As in the case of DNA with •G_N1_C pair, modes of stretching vibration ν(C=O) of guanine and cytosine are shifted into 1788 cm^−1^ and 1760 cm^−1^, as well as scissoring vibration δ(NH_2_) of cytosine (to 1708 cm^−1^). δ(NH_2_) of guanine is absent, due to the lack of hydrogen at N2 site. Stretching vibration ν_sym_(O–P–O) and ν(O–P–O) are shifted into 1070 cm ^−1^ and 853 cm ^−1^, respectively. The averaged O-P-O bonding angle is 97.9° ± 8.7°, whereas the averaged P–O bond length is equal to 1.70 ± 0.31 Å.

**Figure 8.**
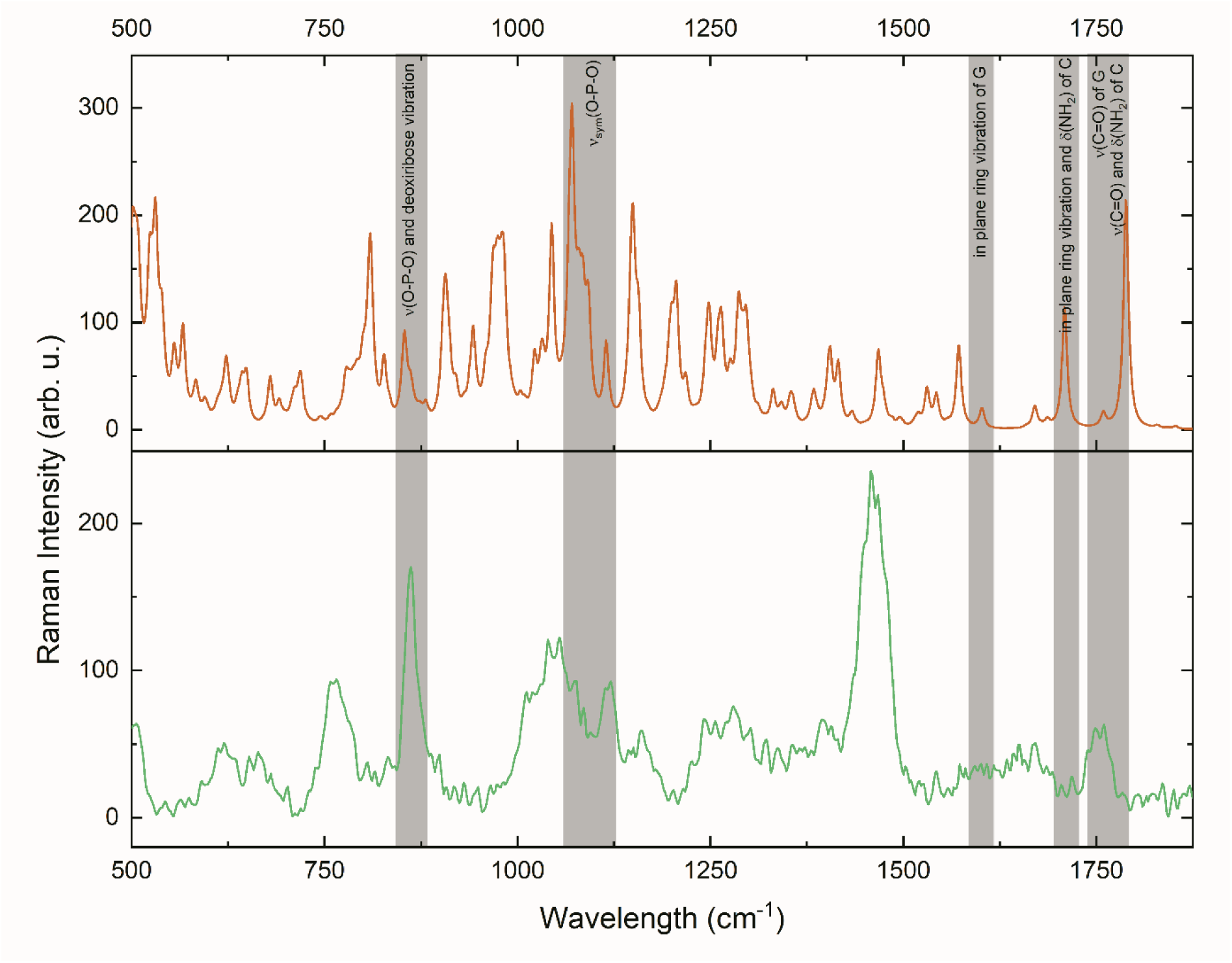
The comparison of spectra of DNA irradiated with the dose of 4000 protons (lime green) with calculated Raman spectrum of damaged DNA with •G_N2_C pair (orange)

**Figure 9.**
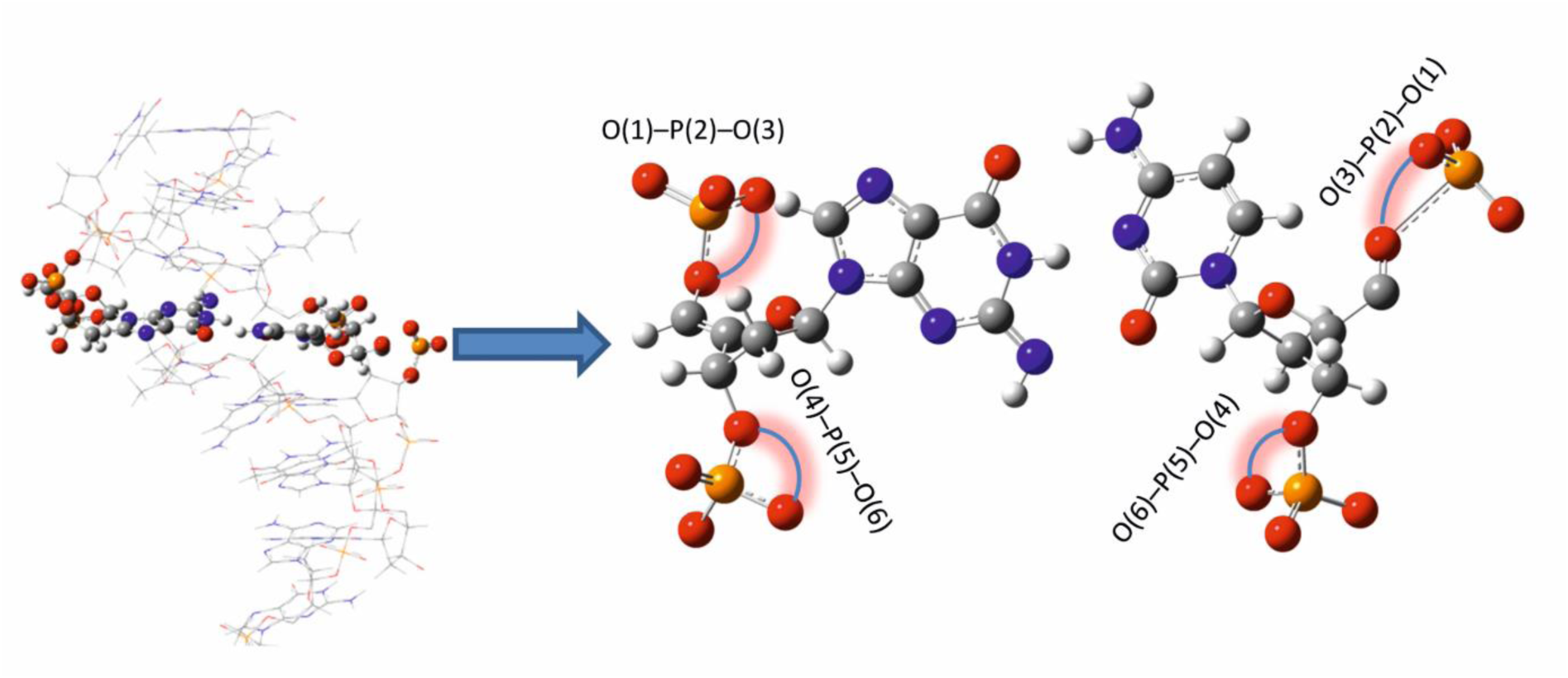
Schematic model of the theoretical molecular system of DNA (left) with •G_N2_C pair (right) treated quantum mechanically

### •G_N1,N2_C

Calculated spectrum of DNA with •G_N2_,_N2_C pair and experimental spectra of irradiated DNA are presented in Figure 10. Figure 11 shows a schematic model of optimized structure. Stretching vibration ν(C=O) of cytosine and guanine and scissoring vibration δ(NH_2_) of cytosine are shifted into 1782 cm^−1^, 1756 cm^−1^, and 1720 cm^−1^, respectively. Symmetrical stretching vibration ν_sym_(O–P–O) is shifted into 1121 cm^−1^. The averaged O–P–O bond angle is in case of this structure 95.3° ± 3.1° and the P–O bond length is equal to 1.80 ± 0.33 Å.

**Figure 10.**
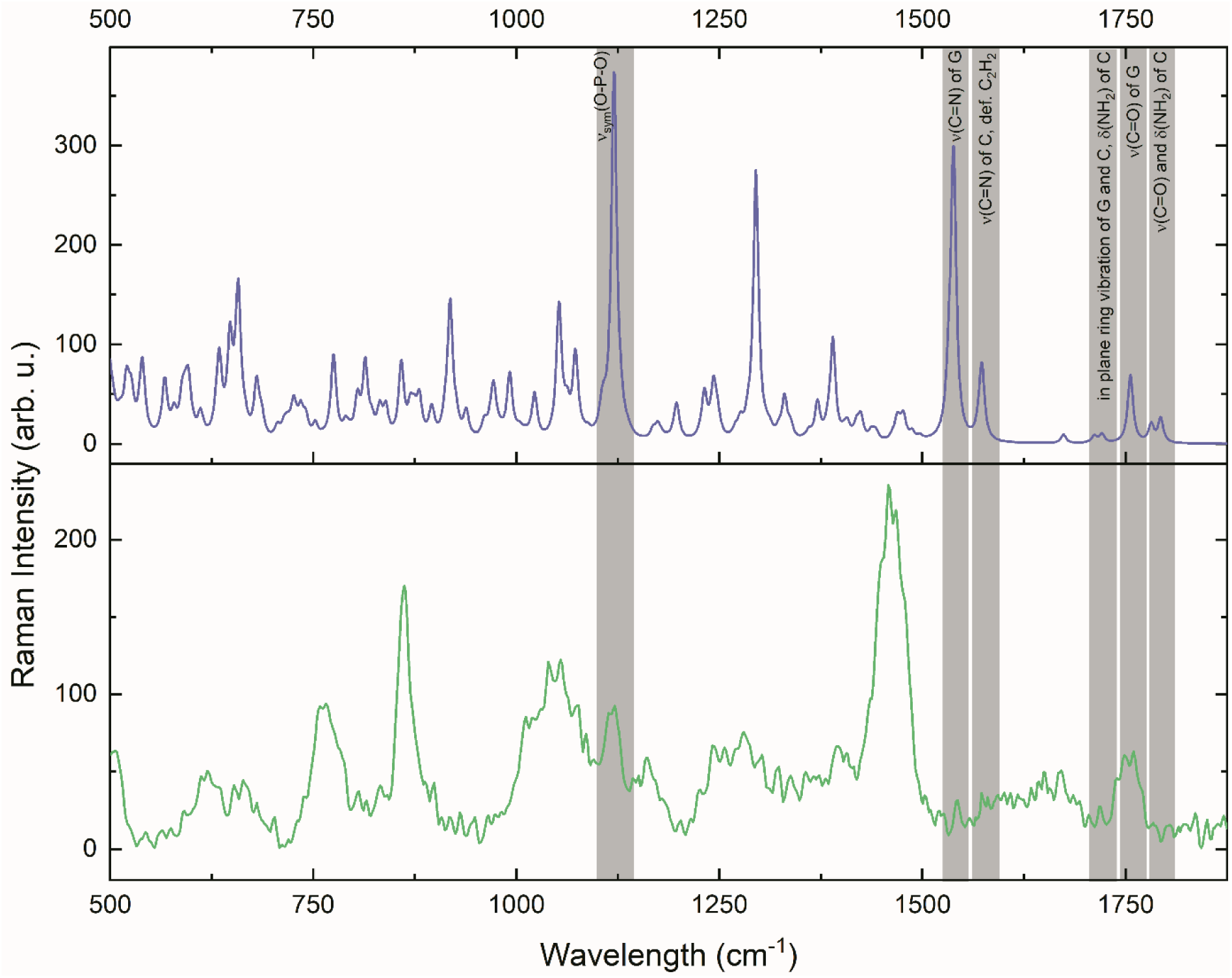
The comparison of spectra of DNA irradiated with the dose of 4000 protons (lime green) with calculated Raman spectrum of damaged DNA with •G_N1_,_N2_C pair (lilac)

**Figure 11.**
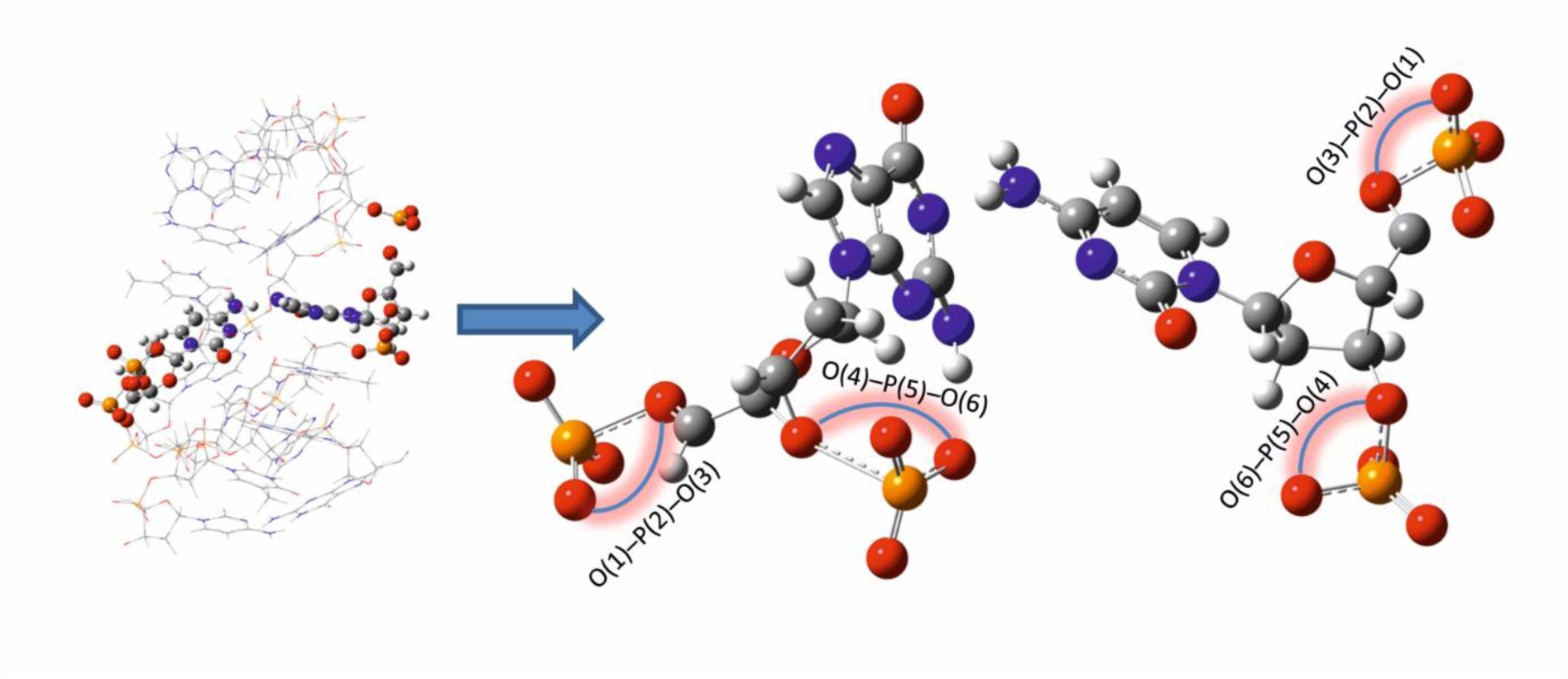
Schematic model of the theoretical molecular system of DNA (left) with •G_N1_,_N2_C pair (right) treated quantum mechanically

A direct comparison of experimental and theoretical spectra of control (not irradiated) DNA presented in Figure 2 proves proper selection of the theoretical molecular system as well as calculation procedure. High correlation between experimental and theoretical results is visible at first glance. Each marker band of DNA bases and the backbone, listed in results section, was observed in theoretical spectrum.

Proposed modifications of molecular system used for calculation allowed to explain spectral differences between experimental spectra of control and irradiated DNA (Figure 4). Our calculations show that the lack of the proton between the bases and consequently the break of hydrogen bond between these bases significantly changes the DNA structure. In the case of all simulated DNA damage, the presence of neutral guanine radical
 in guanine–cytosine pair results in an increase of the length of the P–O bond in phosphate groups, while the O–P–O bonding angle in all optimized damaged structures decreases. As a consequence, the distance between base pairs is smaller than in the case of unchanged DNA, which suggests a local change of DNA conformation.

Observed in experimental spectra peak split and shift of the band νsym(O–P–O) from 1069 cm^−1^ into 1017 cm^−1^, hence the change in potential of part of O–P–O bonds, also indicates that some amount of the DNA changes its conformation. This effect was observed in spectra calculated for DNA with •G_N2_C and •G_N2_,_N2_C pairs. It is known that changes of band intensity of ν(O-P-O) at 1240 cm^−1^, 1190 cm^−1^, and 1080 cm^−1^ in the DNA backbone are associated to strand breaks, whereas ν(C-O) at 1060 cm^−1^ and 880 cm^−1^ to the structure change of sugar moiety (40).

Moreover, for calculations for DNA with •G_N2_C and •G_N2_,_N2_C pairs we have observed an increase of P–O bond lengths to the level above 2 Å for one bond in case of •G_N2_C and two bonds in case of •G_N2_,_N2_C, while P-O bonds in phosphate esters range within 1.59–1.63 Å. This extreme increase of bond lengths is related to the phosphodiester cleavage and the break of the DNA backbone (41).

The break of the hydrogen bond between the bases in the GC pair causes the rotation of these bases with respect to each other for all optimized structures of damaged DNA. This, in turn, prevents base stacking, one of the phenomena maintaining DNA. Such DNA structure is not stable anymore, which can explain the break of P–O bonds in the DNA backbone. The shift of the so called base stacking mode ν(C=O) can also be explained by the unstacking of bases caused by proton exposure (40), which can be seen in experimental as well as calculated spectra for all the three models of damaged DNA. This result shows that even one single local change in bonding between bases can result in the change of DNA conformation and may lead to the DNA strand break.

Our previous work indicated that the bond between carbon and oxygen from the DNA backbone is the most sensitive for the photon exposure and it breaks upon interaction with •H and •OH radicals (24). In contrast to our previous finding, in this article we prove that the interaction with guanine radicals may lead to the break of different P-O bonds included in the DNA backbone. Such radiation damage can be observed in Raman spectra. Raman markers of this radiation damage include i) an intensity decrease of the bands related with νsym(O-P-O) and νasym(O-P-O) and their shift towards higher energy (lower wavenumber) as presented in Table 1, ii) the appearance of ν(C=O) and δ(NH2) from cytosine, base stacking vibration, and ν(C=O) and δ(NH_2_) from guanine.

The same effect was observed by Zhu et al. in 2008. Using Raman spectroscopy the authors studied calf thymus DNA and nucleotides in aqueous solution after 9 min, 20 min, and 40 min ultraviolet radiation exposure. The results proved that the influence of ultraviolet radiation is stronger on single nucleotides than on the whole DNA molecule. However, the authors proved that the molecular conformation of the DNA was changed and the hydrogen bonds were damaged. The researchers also described the interaction of UVR with purines and pyrimidines, which were badly damaged after exposure (42). Structural changes of double stranded DNA in aqueous solution induced by γ radiation were studied by Fourier-Transform-Raman spectroscopy by Sailer et al. The authors put forward that intensity of the bands associated to T, A, G, C increases, which indicates unstacking of these bases (40). The same effect is observed in the presented research at 1335 cm^−1^ and ν(C-N) at 1306 cm^−1^ in C and G (32).

## Conclusions

In conclusion, we have demonstrated that the presence of a guanine radical in DNA double strand and hence the interaction disturbance between the nitrogen bases, which can be induced by proton radiation, has a significant effect on the DNA structure. This includes the change of DNA conformation as well as single or double break of DNA strand. High sensitivity of Raman spectroscopy allowed for detection of DNA radiation damage caused by exposure to 2 MeV protons. Spectral changes related to radiation exposure were explained on the basis of theoretical DFT calculations. The combination of theoretical and experimental data proved that a small change in DNA bases such as oxidation of nitrogen destabilizes the DNA structure locally. Due to the weaker interaction between bases, the DNA backbone may locally change its conformation. These conformational changes determine DNA properties (43) and can affect the radiation sensitivity. Therefore, such results are of significant importance for radiation research.

## Acknowledgments

This research project has been financed by the funds from the National Science Centre (Poland) granted on the basis of decision no. 2014/13/D/NZ1/01014. This research was supported in part by PL–Grid Infrastructure. Mr. Zbigniew Szklarz and Mr. Tomasz Pieprzyca are gratefully acknowledged for help in sample irradiation.

